# FLInt: Single Shot Safe Harbor Transgene Integration via *F*luorescent *L*andmark *Int*erference

**DOI:** 10.1101/2023.01.19.524728

**Authors:** Nawaphat Malaiwong, Montserrat Porta-de-la-Riva, Michael Krieg

## Abstract

The stable incorporation of transgenes and recombinant DNA material into the host genome is a bottleneck in many bioengineering applications. Due to the low efficiency, identifying the transgenic animals is often a needle in the haystack. Thus, optimal conditions require efficient screening procedures, but also known and safe landing sites that do not interfere with host expression, low input material and strong expression from the new locus. Here, we leverage an existing library of *≈* 300 different loci coding for fluorescent markers that are distributed over all 6 chromosomes in *Caenorhabditis elegans* as safe harbors for versatile transgene integration sites using CRISPR/Cas9. We demonstrated that a single crRNA was sufficient for cleavage of the target region and integration of the transgene of interest, which can be easily followed by loss of the fluorescent marker. The same loci can also be used for extrachromosomal landing sites and as co-CRISPR markers without affecting body morphology or animal behavior. Thus, our method overcomes the uncertainty of transgene location during random mutagenesis, facilitates easy screening through fluorescence interference and can be used as co-CRISPR markers without further influence in phenotypes.

## Introduction

The ability to engineer transgenic and mutant animals has afforded one of the biggest revolutions in life sciences. *Caenorhabditis elegans* is a popular laboratory animal, with ten thousand strains carrying exogenous, recombinant DNA available. The first transgenic *C. elegans* animals were generated by microinjection into the worm’s gonad to establish extrachromosomal arrays (Stinchcomb et al. 1985). These arrays are, however, unstable, do not follow Mendelian inheritance and get lost mitotically, leading to mosaic animals in which not all somatic cell express the transgene. When the ectopic DNA is not accompanied by a visible marker, this effect can be misinterpreted as a lack of phenotype. Several strategies have been proposed to circumvent this phenomenon, from the enrichment of the transgenic animals using antibiotic selection (Semple et al. 2010; Giordano-Santini et al. 2010; Radman et al. 2013) to rescue from strong phenotypes such as temperature-sensitive lethality (*pha-1(ts)*) (Granato *et al*. 1994) or paralysis (*unc-119*) (Maduro 2015), however, none of them succeeded in eliminating the mosaic expression. Furthermore, extrachromosomal arrays contain large copy numbers of the injected DNA, which often causes overexpression artefacts, but have the advantage that transgenes become visible even beyond their native levels. For example, many fluorescent tags to endogenous proteins are poorly visible due to their low expression levels and promoter activity (Das et al. 2021; Walker et al. 2000). The problem of unstable inheritance can be mitigated by integrating the transgenic array. Traditional integration methods are based on random mutagenesis, either using a gene gun (Praitis et al. 2001), that allows integration at low frequencies, or chemicals like UV/TMP, X-ray irradiation (Mariol *et al*. 2013) or singlet oxygen generators (miniSOG) (Noma and Jin 2018). However, cumbersome and time-consuming screening efforts are necessary to identify the integrants, and the locus of integration remains unknown unless subsequent mapping experiments are conducted. In addition, the mutagenesis causes extensive DNA double strand breaks, and thus, the resultant animals needs to be backcrossed several times and verified to ensure minimal genetic variability. Even though targeted, MOS-transposase directed, single copy integrations (Frøkjær-Jensen *et al*. 2008, 2012), recombination-mediated cassette exchange (Nonet 2020, 2021) and CRISPR transgenesis (Friedland *et al*. 2013; Paix *et al*. 2017; Dickinson *et al*. 2015) are available, extra-chromosomal arrays were and still are the standard in many laboratories for fast and efficient generation and screening of transgenic phenotypes.

Over the last few years, many different methods have been proposed and demonstrated for site-directed CRISPR/Cas9 mediated locus-specific integration of ectopic DNA such as extra-chromosomal arrays (Yoshina *et al*. 2016; El Mouridi *et al*. 2022) or single copy transgenes (Silva-García *et al*. 2019; El Mouridi *et al*. 2022) into safe habor integration sites. These methods rely on a crRNA that recognizes a single site in the genome and facilitates Cas9 mediated double strand DNA breaks. The subsequent non-homologous end joining or homology-directed repair probabilistically integrates the co-delivered ectopic DNA. Even though these methods overcome many of the above-mentioned shortages of unstable transgenesis and variable expression, so far there are only a limited number of target sites available (e.g. *ben-1, dpy-3*, MosSCI) (Yoshina *et al*. 2016; Frøkjær-Jensen *et al*. 2008; El Mouridi *et al*. 2022). Recently, Frokjaer-Jensen and colleagues generated a library containing 147 strains carrying single-copy loci expressing the red fluorophore tdTomato in somatic nuclei, in addition to 142 nuclearly localized GFP strains (Frøkjær-Jensen et al. 2014), which have aided mapping and in genetic experiments (Das et al. 2021; Fay 2006; LaBella et al. 2020; Noble et al. 2020). Originally, these strains were generated as dominant genetic markers and can also be used as landmarks to map genetic position of mutants and transgenes. Because the integrated transgenes of many of these strains locate to intergenic regions and are transcriptionally active, we reasoned that these loci would satisfy many if not all conditions as further safe-harbor integration sites.

Here we leverage these strains and demonstrate that a single crRNA can cut the tdTomato DNA sequence at extremely high efficiency, affording a selection of 147 possible integration sites, 121 of which are intergenic Frøkjær-Jensen *et al*. (2014). Moreover, the loss of tdTomato fluorescence during the integration not only facilitates screening purposes, but can also be used as co-CRISPR marker during gene-editing at distant loci. Importantly, we show that the integration of a model transgene per se does not affect worm physiology, and even intragenic insertions appear to be phenotypically silent. This method has considerable advantages in multiplexed genome engineering, when the co-CRISPR locus cannot be unlinked easily from the editing site. Lastly, we demonstrate the potential of the single copy GFP sites as dominant co-CRISPR marker and homologous repair events identifier through genetic conversion of GFP to BFP with a single nucleotide change.

## Materials and methods

### Animal maintenance

Nematodes were cultivated on NGM plates seeded with *E. coli* OP50 bacteria using standard protocols (Stiernagle 2006; Porta-de-la Riva et al. 2012). All transgenic strains in this study are listed in the Supplementary Table S1. The parental strains carrying *eft-3*p::tdTomato::H2B and *eft-3*p::gfp::H2B used as the identified landing sites from miniMos ((Frøkjær-Jensen *et al*. 2014)) were maintained and cultured at 20°C prior to injection. All strains used in this study can be assessed in Supp Table 2.

### Molecular biology

Gibson assembly was regularly used for plasmid construction. Briefly, specific primers were designed and PCR was performed using KOD DNA polymerase (Sigma Aldrich). The amplification of DNA fragments was done following manufacturer’s instructions into a Bioer GeneExplorer thermal Cycler. The visualization of DNA fragments was done using an Azure c600 (Azure Biosystems) gel imaging device. Gibson assembly was performed by mixing fragments of the different DNAs at a 3:1 ratio (insert:backbone) and a 2X homemade Gibson Assembly Master Mix. The bacterial transformation was done using either NEB® 5-alpha or 5-alpha F’Iq Competent *E. coli*.

The plasmids (Supplementary Table S3) used as the co-injection markers are pCFJ90 (*myo-2*p::mCherry), pCFJ68 (*unc-122*p::GFP) and pCFJ104 (*myo-3*p::mCherry). The plasmids used as the transgene for integration are pNM5 (*nlp-12*p::ChRmine), pNM10 (*cct- 2*p::mtagBFP2::myosin::spectrin::cryolig2::wrmScarlet(1-10), pNM11 (*mec-4*p::*trp-4*::wrmScarlet), pNM12 (*mec-4*p::RGECO1 syntron), pNM13 (*ges-1*p::CRE), pNM14 (*rab-3*p::CRE), pNMSB91 (15xUAS::*delta pes-10*p::ACR1), and pHW393 (*rab- 3*p::GAL4). The injection mix was prepared by mixing the plasmid of interest, the co-injection markers, and DNA ladder (1 kb Plus DNA Ladder, Invitrogen) at varying ratios. All primer sequences are available in Supp Table 4.

### crRNA design and selection of the target sequence

All crRNAs were designed using Benchling’s DNA editor with single guide option, 20-nt length, PAM sequence (NGG) and were purchased from Integrated DNA Technologies (IDT, Sup, Fig. 5). The crRNA against tdTomato (5’-GTGATGAACTTCGAGGACGG|CGG-3’) recognizes two sites in the tdTomato gene due to the tandem repeat (Fig. 1). The recognition sites are at the 306th and the 1032th nucleotides. Off and on-target specificity has been compiled with CRISPOR (Concordet and Haeussler 2018). Off-target sites that are recognized with 4 mismatches include *ubc-3, gcy-11, Y73F8A*.*5, C55B7*.*3* and *F10G8*.*1*. The crRNAs against gfp excise DNA at the middle of the gene (5’-CTTGTCACTACTTTCTGTTA-3’) and 3’ down-stream region (5’-TGAACTATACAAATGCCCGG-3’). All HR template sequences are shown in Supp Table 6.

**Figure 1.**
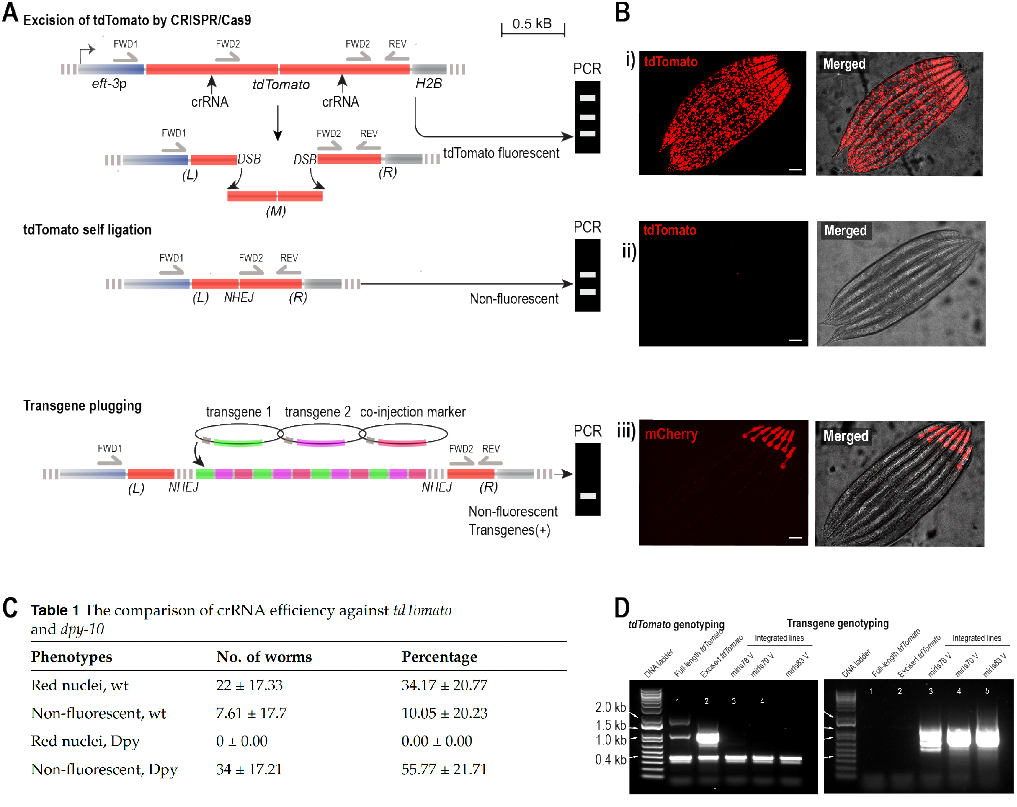
Principle and expected outcomes of FLInt **A, B:** A) Sketch of the different genetic interventions and B) expected phenotypic outcome in tdTomato or transgene fluorescence. (i) Red nuclear fluorescence indicating parental tdTomato fluorescence; (ii) Loss of red indicates successful gene editing; (iii) target transgene fluorescence and loss of red nuclear fluorescence as candidates for stable transgenesis. **C:** Table with the crRNA cleavage efficiency at the tdTomato locus at *oxTi553* compared to the well characterized *dpy-10* locus. Similar efficiencies have been found for tdTomato sites distributed over all 6 linkage groups (see also Fig. 3). **D:** A three-primer PCR genotyping strategy can be used to follow landing site disruption and successful integration. Primers are designed to reveal expected bands for an unedited, edited and integrated tdTomato locus (A).

### Off-target assessment of the crRNA

We assessed off-target gene editing of the loci mentioned in the previous section. With the off-target analysis using CRISPOR(Concordet and Haeussler 2018), we selected a candidate gene, C55B7.3 (I:1.17 +/-0.000 cM), for verifying whether it could be recognized and edited while integrating the transgenes on the tdTomato locus. The C55B7.3 gene was amplified from the integrated strains generated by tdTomato excision. Ten animals were pooled from 15 strains (MSB1110, MSB1111, MSB1112, MSB1113, MSB1115, MSB1116, MSB1117, MSB1118, MSB1119, MSB1120, MSB1121, MSB1122, MSB1123, MSB1124, MSB1125). The lysates were prepared using a variation of the single worm DNA extraction described in (Williams *et al*. 1992). Briefly, 10X PCR buffer from BIOTAQ™ DNA Polymerase (Bioline, Cat. No. BIO-21040) was diluted to 1X and supplemented with proteinase K (Fisher Scientific, Cat. No. 10181030) at 0.1*µ*g/*µ*l final concentration. Each worm was lysed in 10*µ*l lysis buffer and incubated at 65°C for 10 min and 95°C for 2 min in a thermal cycler. 90 *µ*l of milliQ water were added to the lysis reaction and 1 *µ*l used as template for PCR. The PCR primers were designed by CRISPOR; forward primer (5’-TCGTCGGCAGCGTCCTTCCCGAGCAAGAAGGGTG-3’) and reverse primer (5’-GTCTCGTGGGCTCGGTGGAACTTACCGTCACCGAAG-3’). The PCR amplicons were sequenced using the 5’-CTTCCCGAGCAAGAAGGGTG-3’ primer. The off-target effect was assessed by comparing the sequencing data to the wildtype nucleotide sequence.

### Microinjection

Similar to the preparation of the conventional injection mix (transgene DNA + co-injection markers, Supp Fig. 7) (Rieckher and Tavernarakis 2017), this method requires an additional portion of CRISPR reagents. The CRISPR mix was prepared by mixing 14 µM of crRNA, 14 µM of Alt-R® CRISPR-Cas9 tracrRNA (IDT), and milliQ water. The crRNA-tracrRNA dimer was induced by incubating the mix at 95°C for 5 min and RT for 5 min. Then, *Streptococus pyogenes* Cas9 nuclease (IDT) was added to form the RNP complex. The CRISPR mix was aliquoted into PCR tubes (2 µL each) and stored at -20°C for further use. The injection mix was prepared by mixing the purified plasmid DNA (Zymo D4016 PLASMID MINIPREP-CLASSIC) with DNA ladder (1 kb Plus DNA Ladder, Invitrogen), 100 ng/µl DNA in total (see Supplemetary Table S1). We added the 2 µL of CRISPR mix (mentioned above) into the 8 µL injection solution to make a total of 10 µl. The mix was centrifuged at the highest speed for 8-10 minutes before injecting. The transgenic strains used as the P0 animals were established by miniMos technique (Frøkjær-Jensen et al. 2014) expressing tdTomato and GFP in all cellular nuclei. We selected the following transgenic strains:

EG7835 [*oxTi556* I (*eft-3*p::tdTomato::H2B)],

EG7846 [*oxTi700* I (*eft-3*p::tdTomato::H2B)],

EG7860 [*oxTi677* II (*eft-3*p::tdTomato::H2B)],

EG7866 [*oxTi564* II (*eft-3*p::tdTomato::H2B)],

EG7898 [*oxTi619* III (*eft-3*p::tdTomato::H2B)],

EG7900 [*oxTi546* III (*eft-3*p::tdTomato::H2B)],

EG7905 [*oxTi390* IV (*eft-3*p::tdTomato::H2B)],

EG7911 [*oxTi705* IV (*eft-3*p::tdTomato::H2B)],

EG7944 [*oxTi553* V (*eft-3*p::tdTomato::H2B)],

EG7945 [*oxTi543* V (*eft-3*p::tdTomato::H2B)],

EG7985 [*oxTi566* X (*eft-3*p::tdTomato::H2B)],

EG7989 [*oxTi668* X (*eft-3*p::tdTomato::H2B)],

EG8958 [*oxTi1022* I (*eft-3*p::gfp::NLS)], and

EG8888 [*oxTi936* X (*eft-3*p::gfp::NLS)]. All transgenic animals that we used as the background strains are available in CGC.

### Visual screening of transgenic animals

The screening of the fluorescent progenies from P0 was performed using a fluorescent stereomicroscope (SMZ25, Nikon Instruments) equipped with a white-light LED light source (Lumencor, Sola S2). We searched for the non-red animals with co-injection marker expression, called positive F1, 3-day post injection. Then, we singled them out into new NGM/OP50 plates. The individual positive F1 were cultured for 3 days at 25°C, and plates were searched for F2 progenies with high transmission frequency (approx. 75%). Six F2 of each of those plates were singled out. After 3 day, the F3 progenies were checked for homozygous expression of the co-injection marker and, if integration had taken place, the integrated lines were characterized. The F3 progenies from the same F1 are determined as identical transgenic line. We calculated the integration efficiency by (no. of integrated line / no. of positive F1) x 100.

### Determination of the integration efficiency on different loci

Six different tdTomato landing sites in different chromosomes were used for assessing integration effeciency: EG7846 (*oxTi700*, I:22.30), EG7860 (*oxTi677*, II:-12.17), EG7900 (*oxTi546*, III:11.80), EG7905 (*oxTi390*, IV:-26.93), EG7944 (*oxTi553*, V:0.29), and EG7985 (*oxTi566*, X:-4.88) (Fig. 3A). Animals were injected with 2 ng/µL, 98 ng/µL DNA ladder (Invitrogen), and tdTomato CRISPR mix. Unless otherwise specified, the P0 animals were cultured at 25°C after injection, as well as the F1, F2, and F3. The integration efficiency was then calculated from three experimental replicates.

### Integrated copy number analysis with qPCR

qPCR was used for detecting and measuring the copy number of the integrated pCFJ90 (*myo-2*p::mCherry) of nine integrated strains (MSB884, MSB886, MSB898, MSB905, MSB911, MSB912, MSB913, MSB914, and MSB915). Sample preparation was done by culturing worms in peptone-enriched plates with NA22 as food source. When plates were full of adult worms, they were washed off the plates with M9 buffer, excess bacteria eliminated by successive washes and lysed in 500 *µ*l lysis buffer supplemented with proteinase k (see *Off-target assessment of the crRNA* section above). The genomic DNA was purified using the Zymoclean Gel DNA Recovery Kit (Zymo Research). qPCR analyses were carried out by AllGenetics & Biology SL (www.allgenetics.eu). Briefly, absolute qPCR was performed with primers indicated in table S3. The qPCR experiment was performed in triplicate for each sample and controls. The qPCRs reactions were carried out in a final volume of 20 µL, containing 10 µL of NZY qPCR Green Master Mix ROX plus (NZYTech), 0.4 µM of the amplification primers, 2 µL of template cDNA, and ultrapure water up to 20 µL. The reaction mixture was incubated as follows: an initial incubation at 95 ºC for 10 min, followed by 40 cycles of denaturation at 95 ºC for 15 s, annealing/extension at 65 ºC for 1 min. A five point 10-fold serial dilution of a known number of copies of the genes under study was used to establish the standard curve and evaluate the reaction efficiency. These dilutions were also performed in triplicate. The Y-intercept and slope were also obtained from the standard curve. Copy number was calculated by the formula: copy number = 10^(Cq - Yintercept)/(slope)^. Copy number of integrated transgenes was obtained by normalizing with *rps-25*.

### Screening for loss of tdTomato fluorescence as a ‘coinjection’ marker

Having multiple transgenes or multicolour phenotype could negatively affect animal health as it constitutes a metabolic burden and limits the degrees of experimental freedom during microscopy experiments (e.g. multicolor imaging acquisitions). Importantly, the above mentioned integration protocol and simplicity of the screening procedure also facilitates the integration of transgenes without the use of visible markers, e.g. such as the PHA::mCherry. To demonstrate this, we generated a dual-fluorescence CRE/lox reporter strain (based on SV2049) with constitutive BFP expression and conditional, CRE-dependent mCherry expression, with the ubiquitous tdTomato expression from the landing site in the background (MSB934). After injecting this strain with a plasmid encoding for an intestinal CRE (*ges-1*p::CRE) together with tdTomato CRISPR mix, we confirmed loss of tdTomato and a BFP/mCherry colorswitch in intestinal nuclei in the F1. Importantly, the intestinal red fluorescence is indicative for the tissue specific CRE-recombination, that would otherwise be obscured had the tdTomato cleavage not taken place. To isolate homozygous integrants, we followed the CRE-dependent BFP/mCherry color switch during the F3 (Fig. S4). We also demonstrated the co-injection marker free integration using the binary UAS/GAL4 expression system (Wang *et al*. 2017), and integrated a panneuronal *rab-3*p::cGAL4 driver construct in the background of a silent UAS::GFP effector strain carrying the tdTomato landing site. Following our experimental pipeline, we obtained positive F1 that panneuronally expressed GFP signal with the loss of tdTomato (Fig. S4). Our results demonstrate that the negative selection due to fluorescent interference of the tdTomato landing site facilitates the screening step in *C. elegans* transgenesis and serves as a safe harbor for transgene expression.

### Integration of extrachromosomal array using FLInt

The integration of the existing extrachromosomal array was done first by crossing the strain of interest to the desired tdTomato marker strain. A CRISPR injection mix containing 14 µM of crRNA against tdTomato, 14 µM crRNA against Ampicilin resistance gene (AmpR), 28 µM of tracrRNA and Cas9 endonuclease was injected in the resulting strain and the progeny scored for loss of tdTomato expression. 100% transmission of the extrachromosomal marker was used as an indicator for integration.

### Screening integrations with PCR

To follow the double-strand break, excision and integration efficiency at the tdTomato site, we designed PCR primers that bind to several regions along the tdTomato gene (Fig. 1A); (1) A forward primer that binds to the region upstream the tdTomato gene, in the *eft-3* promoter (FWD1: 5’-TTTATAATGAGGTCAAACATTCAGTCCCAGCGTTTT-3’) (2) another forward primer that binds to the middle of the gene in both tandem repeats, downstream the excision sites (FWD2: 5’-GACCCAGGACTCCTCCCT-3’), (3) the reverse primer, that binds at the end of tdTomato ORF (REV: 5’-TTACTTGTACAGCTCGTCCATGC-3’). This strategy gives rise to 3 bands when genotyping tdTomato (Fig 1A,C). We utilized this technique for investigating the tdTomato gene before and after being excised by CRISPR/Cas9. The full-length tdTomato is recognized by the 4 binding sites of the 3 primers amplifying three different band sizes: 1.7 kb, 1.1 kb, and 0.4 kb (Fig. 1D, lane 1). The excised tdTomato splits the middle chunk of gene, losing one primer binding site. Only two PCR bands (1.1 kb and 0.4 kb) were detected (Fig. 1D, lane 2). Lastly, in integrated strains only the smallest band (0.4 kb), outside of the integration region is amplified (Fig. 1D, lanes 3-5). To avoid competition between the two different FWD primers, the following PCR conditions proved optimal: FWD primer (1) = 2mM; FWD primer (2) = 0.2mM; REV primer = 2mM; Tm = 55°C; Extension time = 1 min.

### Screening for *lat-1::loxP::*Δ*mCherry* insertions using td-Tomato as Co-CRISPR

The insertion of a loxP site into *lat-1* locus was done using CRISPR/Cas9. To excise *lat-1* gene, we introduced the crRNA (5’-ATGTACACGCATCAAAGATA-3’) (IDT), tracrRNA (IDT), and Cas9 (IDT). The loxP site and additional sequence (Δ*mCherry*) insertion and PAM mutation was induced by the HR template (Table Sx) with 35-nt homology arms (IDT). The CRISPR mix was prepared followed the details above and injected into the gonad of the background strain EG7944 (*oxTi553* V [*eft-3*p::tdTomato::H2B]). The concentration of the homology repair template was 167ng/ul. The screening of F1 was done after 3 days using the fluorescent microscope. The candidate F1(s) were selected from the jackpot plates based on the loss of tdTomato fluorescent signals among the F1 population. The candidates were singled out onto new NGM/OP50 plates before genotyping. To genotype the loxP insertion, worms were lysed and genotyped as detailed in the *Off-target assessment of the crRNA* section with primers 5’-CGATGTTGACAACTGAAGTGA-3’ and 5’-GGTAATTTCTGACATGGCTCA-3’. The edits were observed in an electrophoresis gel by the shift of the edited DNA band (417 bp) compared to the wildtype (291 bp). The efficiency of *lat-1::loxP::*Δ*mCherry* insertion from each jackpot plate was calculated by (no. of edits / no. of candidate F1) x 100.

### Screening of GFP color switch as the HDR-mediated co-CRISPR marker

The HDR-mediated fluorescent conversion from GFP to BFP (P4) was done with the *eft-3*p::GFP::NLS background strains, EG8888 [*oxTi936* X] and EG8958 [*oxTi1022* I]. The single point mutation of *gfp* gene was triggered by DNA double-strand break via CRISPR/Cas9 approach followed by the HDR that introduces the change of amino acid from the background (Y66H). To do this, the crRNA against *gfp* (5’-CTTGTCACTACTTTCTGTTA-3’), tracrRNA, Cas9 nuclease, and the HR template (5’-TTAAATTTTCAGCCAACACTTGTCACTACTTTCTGTTATGGTGTTCAATGCTTCTCGAGATACCCAGATCATAT-3’; see Supp. Table 6), purchased from IDT, were injected into the P0 animals. After 3-day post injection, the F1(s) progenies were screened for the loss of GFP single which replaced by the expression of BFP in the nuclei. The candidates were then singled out and screened for few generations to obtain the homozygous genotype.

### Fluorescence microscopy

The fluorescence signal of the worms was observed using a confocal microscope (Andor DragonFly 502, Oxford Instruments) attached to a Nikon Eclipse Ti2 inverted microscope body through either a 20x 0.75 oil or a 60x 1.3 oil immersion lens and a back-illuminated sCMOS camera (Sona, Andor). The tdTomato fluorescence signal was excited with a 561 nm laser beam (power intensity 30 %, exposure time = 200 ms) and the emitted signal transmitted using a 594 nm filter. The GFP fluorescence signal was excited with a 488 nm laser beam (power intensity 60 %, exposure time = 100 ms) and transmitted using 521 nm filter. The mCherry fluorescence signal was excited with a 514 nm laser beam (power intensity 40 %, exposure time = 300 ms) and transmitted through a 594 nm filter. The P4 and BFP fluorescence signals were excited with a 405 nm laser beam (power intensity 40 % and 20 % respectively, exposure time = 400 ms and 200 ms respectively) and transmitted using a 445 nm filter. The fluorescence signal was captured using Z-scan protocol (0.7 step size) through the confocal apparatus (Andor DragonFly).

### Healthspan assessment

The wt (N2), full-length tdTomato (EG7944), excised tdTomato (MSB910), three *myo-2*p::mCherry integrated lines (MSB1115, MSB1118, and MSB1122) and *myo-2*p::mCherry (extrachromosomal array) animals were cultured and their development, locomotion, body length and lifespan compared (Fig. S3). The fluorescence intensity and development were done in N2, EG7944, MSB910, and MSB1115. Development was assessed based on the worms size over time from L1 to egg-laying adult stage. Synchronized L1 (Porta-de-la Riva *et al*. 2012) were seeded onto NGM/OP50 plates and incubated at 20°C. We captured the worms at L1 stage (prior to seeding), L3 stage (24 hours after seeding), L4 stage (40-48 hours after seeding) and egg-laying stage (72 hours after seeding) based on the developmental time-line of N2 (Porta-de-la Riva *et al*. 2012). We imaged tdTomato fluorescence intensity in young adult worms using the Z scan protocol (step size =1.7 µm) with 20x magnification (20X/0.75 MImm objective lens). The maximum Z-projection was performed using ImageJ (Fiji). Then, the ROI was drawn using segmented line across the body edge. The average intensity was measured and collected from individual worms. On the last day of developmental assessment, the adult animals were placed on the new plate and the moving trace on bacterial lawn captured. The locomotion behavior was observed under the lab-built microscope (WormTracker (Das *et al*. 2021)). We took the sinusoidal wave appearing in the bacterial lawn after the worm passed as reference of the body angle during locomotion.

Body length was captured in a lab-built microscope (Worm-Tracker (Das *et al*. 2021)) using 2x magnification and measured in ImageJ. By using the segmented line tool, the body length was measured from the nose tip to the tail tip.

The lifespan asssay was conducted by counting number of dead and alive worms in FUDR plates until the whole population diminished. The decrease of viability of each strain were plot from as the survival curve. The mean of lifespan was calculated from the average of age from individual animals in each population. Then, the mean of lifespan was compared to wt strain.

### Statistical analysis

No statistical method was applied to predetermine sample size based on data variability. All data sets were first tested for normality using KS test or Tukey adjusted ANOVA for multiple comparisons as indicated in the Figure legends.

## Results and discussion

### Single tdTomato transgenes as safe harbor landing pads for exogenous transgenes

To demonstrate that the single tdTomato transgenes can function as versatile sites to integrate transgenes into the genome of *C. elegans*, we designed a single crRNA against the *tdTomato* ORF (Fig. 1A, see Methods) that is not predicted to have a full length off-target binding probability. Because tdTomato is a tandem-dimer gene of a single fluorophore, the successful Cas9 cleavage will cut twice the DNA, excising a large portion of the gene. The concomitant loss of fluorescence should, in principle, facilitate the screening process, and therefore speed up the identification of successful integrations. Thus, the tdTomato site serves a dual function: a successful co-injection marker and a landing site. We first sought to test whether the selected crRNA cleaves the tdTomato sequence. We reasoned that successful dsDNA break results in a loss in tdTomato fluorescence in the filial generations. Indeed, many animals in the F1 of an injected P0 have already lost their tdTomato fluorescence, which is readily identifiable in a normal fluorescence stereomicroscope (Fig. 1B). Some animals, however, showed a considerably lower fluorescence, indicative for a single edit on one parental chromosome. We also frequently observed a mosaic pattern in the somatic cells of the F1s, possibly due to cleavage after the first cell division. These animals would eventually give rise to non-red animals in the F2 generation according to Mendelian segregation. In jackpot broods, we frequently observe 25% of non-red animals from a single injection. We benchmarked the DNA cleavage efficiency for the tdTomato against the widely used, highly efficient *dpy-10* protospacer (Arribere et al. 2014) and coinjected 2µM for both crRNAs together with recombinant Cas9 (Paix et al. 2017). We then screened for non-red and Dpy animals as a readout for simultaneous cleavage of both DNA strands at the *dpy-10* and tdTomato locus. From the total 13 jackpot broods we screened, we found 34% red, wildtype animals, 56% non red, Dpy animals and 0% red, Dpy worms. Since all Dpy animals had also lost the tdTomato fluorescence in the F1, we reasoned that the crRNA for tdTomato is, at least, as efficient as the highly efficient *dpy-10* protospacer (Arribere et al. 2014; El Mouridi *et al*. 2017). In addition, we found 10% non-red, wild type worms (Fig. 1C, Table 1), suggesting a slightly higher efficiency of the td-Tomato protospacer and making FLInt an extremely well suited candidate method for transgene integration at many potential sites across the genome. Together, these results not only indicate that the selected crRNA for tdTomato efficiently guides Cas9 for subsequent DNA cutting, but also that it does so at a high efficiency, allowing identification of events already in the F1. As a last test for the suitability of FLInt as safe habor sites, we assessed if the tdTomato crRNA causes any unwanted off-target effects. The genotyping of nine different strains at the most likely predicted off-target site (containing 4 mismatches), however, did not identify any further edits. Likewise, we also did not detect gross defects in healthspan and locomotion or any other behavioral phenotypes compared to N2 wildtype animals (Fig. S3). We observed a general suppression of a lifespan defect in the parental *oxTi553* strain, which behave poorly at 25CFrøkjær-Jensen *et al*. (2014). This suggests that the edits are not interfering with the normal physiology of the animal and have nearly wildtype behavior (Fig. S3).

**Table 1.** The comparison of crRNA efficiency against *tdTomato* and *dpy-10*

| Phenotypes | No. of worms | Percentage |
| --- | --- | --- |
| Red nuclei, wt | 22 ± 17.33 | 34.17 ± 20.77 |
| Non-fluorescent, wt | 7.61 ± 17.7 | 10.05 ± 20.23 |
| Red nuclei, Dpy | 0 ± 0.00 | 0.00 ± 0.00 |
| Non-fluorescent, Dpy | 34 ± 17.21 | 55.77 ± 21.71 |

Having established highly-efficient DNA cleavage using the tdTomato crRNA, we proceeded to inject 20 P0 animals with the CRISPR mix and a *myo-2*p::mCherry plasmid as transgene-of-interest (TOI) into the *eft-3*p::tdTomato::H2B V strain (EG7944) (Fig. S2A), following loss of red nuclear fluorescence from the tdTomato and gain of mCherry expression in the pharynx during the filial generations (Fig. S2A). Consistent with our prior observations, we found that some F1 had already lost the strong tdTomato nuclear fluorescence displayed by the P0, an indication of the successful excision of both homologous chromosomes in the first generation after injection. We singled out animals positive for red pharynx (Fig. S2B), noticing that most of the transgenic animals that expressed mCherry had also lost nuclear tdTomato expression. To distinguish between expression from the extrachromosomal array and integrants, we selected six F2 animals from high transmission plates (PHA::mCherry, loss of nuclear tdTomato) and eventually obtained 1-3 integrated lines based on 100% transmission frequency in the F3 from one injection (see also Supplementary Table 1).

Often, the transgene of interest does not lead to a visible phenotype, for example effector or driver strains in bipartite expression systems (Das *et al*. 2021; Porta-de-la Riva *et al*. 2021; Yang and Yuste 2017). To follow the integration of such transgenes, we developed a PCR genotyping strategy (Fig. 1A, D, see Methods) using three primers that target the region around tdTomato for amplification, with different amplicon sizes according to the genetic recombination occurred. We selected animals from the three different populations: td-Tomato::H2B (no excision), non-fluorescent (loss of tdTomato) and non-fluorescent/PHA::mCherry (expectedly tdTomato inserted) to isolate their DNA for genotyping. In the parental strain with tdTomato expression, the three primers would anneal (one of them twice) and three bands of different sizes would be amplified. Expectedly, we found that loss of tdTomato signal in absence of the transgenic marker was genomically accompanied by the loss of the longest DNA band, indicative for a successful Cas9 activity, and repair through non-homologous end joining (NHEJ). However, in PHA:mCherry homozygous animals carrying the successful integration, we were unable to amplify the region flanked by the two crRNA target sites. We reasoned that the region with inserted transgene could not be amplified due to the large size of the multicopy transgene, which could be up to millions bases in length (El Mouridi *et al*. 2022). However, the small band corresponding to the end of the td-Tomato gene and downstream the expected integration site (0.4 kb) was amplified, serving as a positive control for PCR (Fig. 1D, lane 3-5). Taken together, these results established that ectopic transgenes can be integrated by CRISPR using site-specific crRNAs into the tdTomato landing sites as multicopy transgenes with very high efficiency and reliability.

During the expansion of the injected animals we consistently observed different integration efficiency based on the culturing conditions. Similarly to what had been previously described for integrations through the miniMos technique (Frøkjær-Jensen et al. 2014), we hypothesized that the temperature at which the P0 is grown after injection might affect the integration efficiency. To investigate this, we reared the injected P0 at 16°C or 25°C for two days until we screened F1 for positive transgenesis events. The F2 and F3 progenies, however, were invariably raised at 25°C. After obtaining the integrated lines, we found that culturing the P0 at 25°C promoted higher integration efficiency compared to 16°C (Table 2). At this temperature, we obtained 100% success rate, with integrated lines from every injection round (3/3). Conversely, only one integrated line was obtained of the P0 incubated at 16°C (1/4, Table 2). This result is in agreement with previous reports in vertebrates and plants showing that Streptococus Pyogenes Cas9 efficiency is higher at elevated temperatures (Moreno-Mateos et al. 2017; LeBlanc et al. 2018) and suggests that transgene integration is temperature-dependent.

**Table 2.**
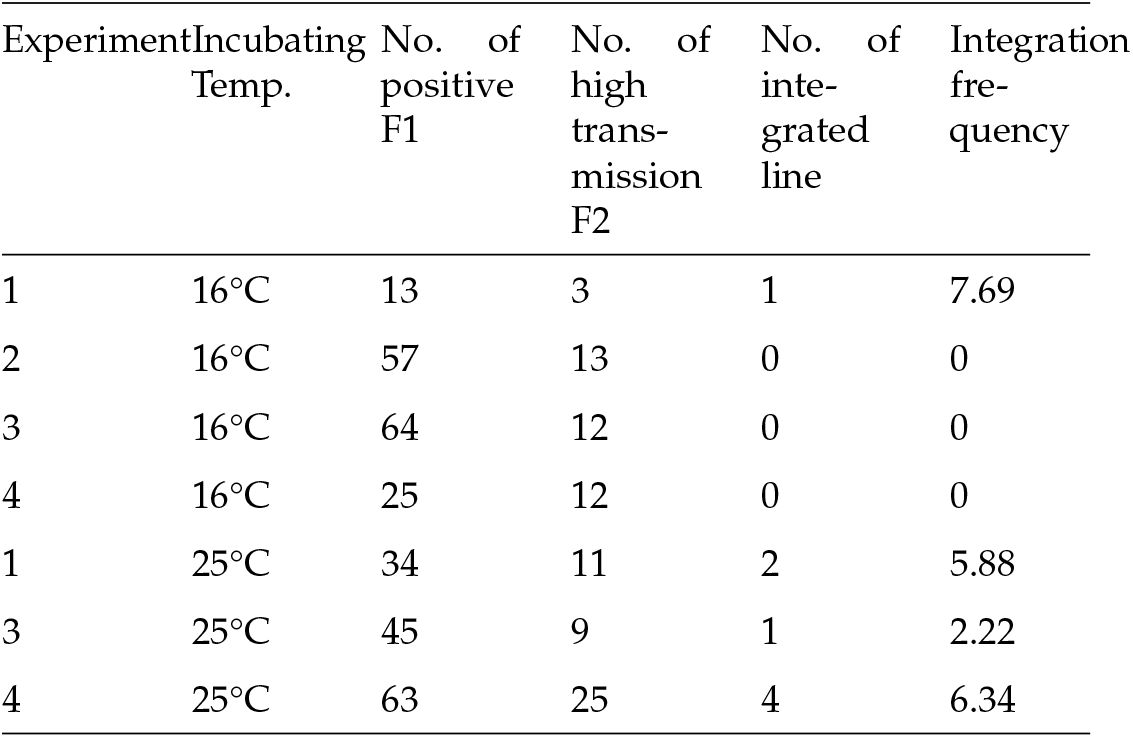
The effect of temperature on FLInt mediated integration EG7944 animals carrying *oxTi553* were injected with *myo-2*p::mCherry and DNA ladder and followed for integration.

| Experiment | Incubating Temp. | No. of positive F1 | No. of high transmission F2 | No. of integrated line | Integration frequency |
| --- | --- | --- | --- | --- | --- |
| 1 | 16°C | 13 | 3 | 1 | 7.69 |
| 2 | 16°C | 57 | 13 | 0 | 0 |
| 3 | 16°C | 64 | 12 | 0 | 0 |
| 4 | 16°C | 25 | 12 | 0 | 0 |
| 1 | 25°C | 34 | 11 | 2 | 5.88 |
| 3 | 25°C | 45 | 9 | 1 | 2.22 |
| 4 | 25°C | 63 | 25 | 4 | 6.34 |

We were then curious to understand how many copies of the coinjected plasmid were integrated into the safe harbor locus and how this related to the relative amount of DNA injected. Previous integration methods suggested a large variability of integrated copies, ranging from few copies (derived from biolistic transformations (Sarov et al. 2012)) to hundreds after integrating traditional extrachromosomal arrays with random mutagenesis (Noma and Jin 2018). We thus injected varying ratio of coinjection marker/transgene together with the tdTomato CRISPR mix into EG7944 *oxTi553* V or EG7846 *oxTi700* and quantified their integrated copy number using quantitative PCR. We found that a higher plasmid ratio led to a higher copy number, which in turn led to a higher transgene expression from the co-injection marker (*myo-2*p:mCherry) (Fig. 2). Thus, a careful titration of injected plasmid would thus facilitate a balanced expression (in our hands ranging from 20 to 150 copies of the transgene) of the desired transgenes in a known safe harbor locus.

**Figure 2.**
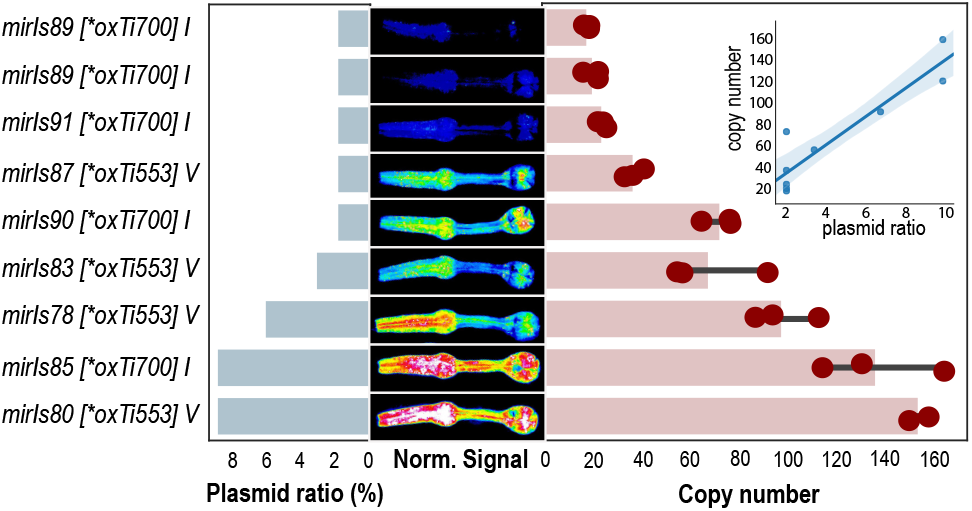
Transgene copy number for nine different transgenes Different relative concentrations of *myo-2*p::mCherry were injected together with the CRISPR mix and a target plasmid (ratio, left bars) and integrated into the same tdTomato locus (oxTi). The resulting homozygous transgene copy numbers were quantified by qPCR (right bars). The middle plot shows the pharynx fluorescence and the inset shows the integrated copy number as a function of the injected plasmid ratio.

**Figure 3.**
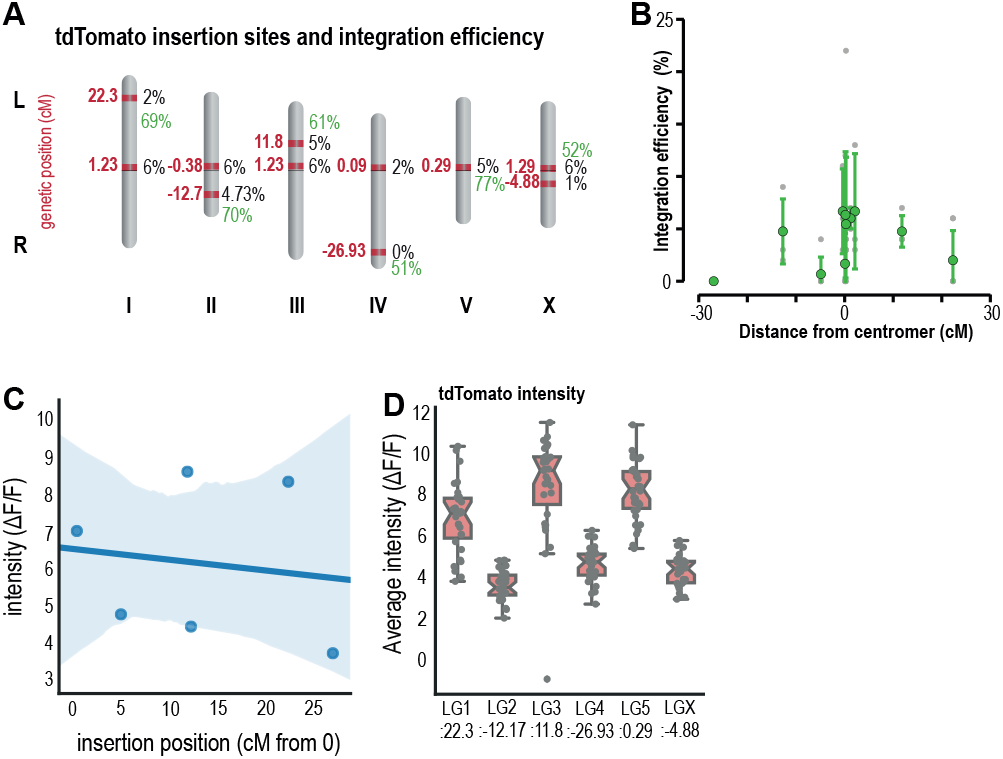
Integration efficiency correlates with chromosomal position **A:** Summary schematic of the different landing sites and their chromosomal position used and their integration/tdTomato cutting efficiencies. The cutting efficiencies to the proximal sites are shown in green. **B:** Plot of integration efficiency vs genetic position irrespective of the linkage group. A strong drop in efficiency is observed for sites close to the chromosomal periphery. Grey are data points from individual experiments, green dots show mean *±* standard deviation for each landing site. See also Supplementary Table 1. **C:** Fluorescence intensity for the single-copy tdTomato transgenes at the indicated sites. **D:** Plot of the tdTomato fluorescence intensity vs chromosomal position.

Taken together, highly efficient integration methods reduce the time consuming screening required in traditional transgene integration procedures. Compared to the conventional method using UV/TMP, in which worms are propagated for several generations during 3-5 weeks before the screening (Mariol et al. 2013) and require posterior outcross to non-mutagenized worms, our method establishes integrated lines within 9 days post injection, essentially bypassing the formation of an extrachromosomal array. Besides, the colorimetric change provides visual, dominant marker for screening, allowing fast identification of positive F1 among the background phenotype. In addition, since the loss of fluorescence only takes place after cleavage of both chromosomes, rapid screening of homozygous edits is facilitated, even granting the omission of another injection marker than the loss of the same tdTomato. For example, we successfully integrated *ges-1*p::CRE and *rab-3*p:GAL4 into their effector strain and recovered transgenic after a single injection (Fig. S4 and Methods). Thanks to previous work in the generation of the many miniMos strains, (Frøkjær-Jensen et al. 2014), a single crRNA can be used on the tdTomato present in single copy in 147 different loci from strains that are available in the CGC, providing high flexibility in designing transgenic animals and downstream experiments. We have named our improved transgene integration method “FLInt” for **F**luorescent **L**andmark **Int**erference, in reference to the locus of transgene incorporation.

### GFP as an alternative FLInt landing site

Often, the choice of the co-injection marker is guided by the transgene of interest. Thus, the tdTomato sites are incompatible if the TOI already contains a tdTomato fluorophore. Likewise, if the transgene encodes for a nuclear localized mCherry, down-stream analysis can be confusing. With the aim of posing an alternative in those cases, We approached the single copy GFP marker strains described in (Frøkjær-Jensen *et al*. 2014) to asses if they could serve as a convenient alternative. We designed a pair of crRNAs to disrupt the GFP ORF (Fig. S5B) generating a deletion and verified that this event led both to a potent loss of GFP expression and to the generation of integrants (Fig. S5B). We then compared the GFP protospacer against the tdTomato protospacer by means of injecting a CRISPR mix that contained these crRNAs into a transgenic animal that had both landing sites. In the screening, we found around 34% of non-red/non-green animals in the F1 and similar frequencies of animals with either loss of green or loss of red (Figure S4B, C), suggesting that both crRNAs have comparable cutting efficiencies if their cognate loci. Together, these demonstrate that the single copy GFP loci serve as good alternative targets for FLInt. However, due to abundant gut autofluorescence and the generally weaker fluorescent signal, transgene screening is more difficult than in the tdTomato strains.

### The efficiency of transgene integration varies with chromosomal position

Having shown that the single copy tdTomato (or GFP) can be used as FLInt landing sites, we wondered if the entire zoo of 147 possible landing sites would accept exogenously delivered transgenes with similar efficiencies. We thus selected a random set of landing sites distributed over all linkage groups and tested the integration potential of *myo-2*p::mCherry into 6 different background strains, one in each chromosome (see Methods) and calculated the integration efficiency from a standardized experiment (same injection mix, different landing sites, see Methods). We found that the most successful and highest efficiency was on chromosome I and II (oxTi556 I, 6.49%; oxTi564 II, 6.46%), followed by the landing sites on chromosomes X and III (oxTi668 X, oxTi619 III), 6.2% and 5.8% respectively (Fig. 3A, Supplementary Table 1). This result showcase that, even when integration is possible on all linkage groups, it may be more probable in some than others, based on causes that are external to the transgene but internal to the landing site of the TOI. Apart of being on different chromosomes, the different landings sites are located on different genetic positions within each linkage group (LG1:22.30, LG2:-12.17, LG3:11.8, LG4:-26.93, LG5:0.29, and LGX:-4.88). In general, we found that the tdTomato landing sites in the center of the chromosome have higher efficiency compared to the farther tdTomato landing sites (Fig 4B). In our hands, only one landing site (LG4:-26.93) was not accepting any transgene integrations after many trials, even though cutting efficiency was comparable to other loci. Taken together, even though integrants can theoretically be obtained on all tested loci (Supplementary Table 1), experimentally, integration frequency varies dramatically between them. Lastly, we asked if the different locations of the insertion sites could possibly lead to differences in downstream transgene expression. Because the integrated DNA exist as multicopy transgene with varying copy number (e.g. Fig. 2), we compared the fluorescence intensity of the tdTomato loci produced by the single copy *eft-3*p::tdTomato::H2B transgene among the six linkage groups used in our experiments (EG7846 I, EG7860 II, EG7900 III, EG7905 IV, EG7944 V, and EG7985 X) and found that the tdTomato intensity from each strain is different (Fig. 3C) but uncorrelated with the genetic position based on their insertion sites. Even though transgene integration at different loci varied, we concluded that transgene expression seemed unaffected (Fig. 3D).

**Figure 4.**
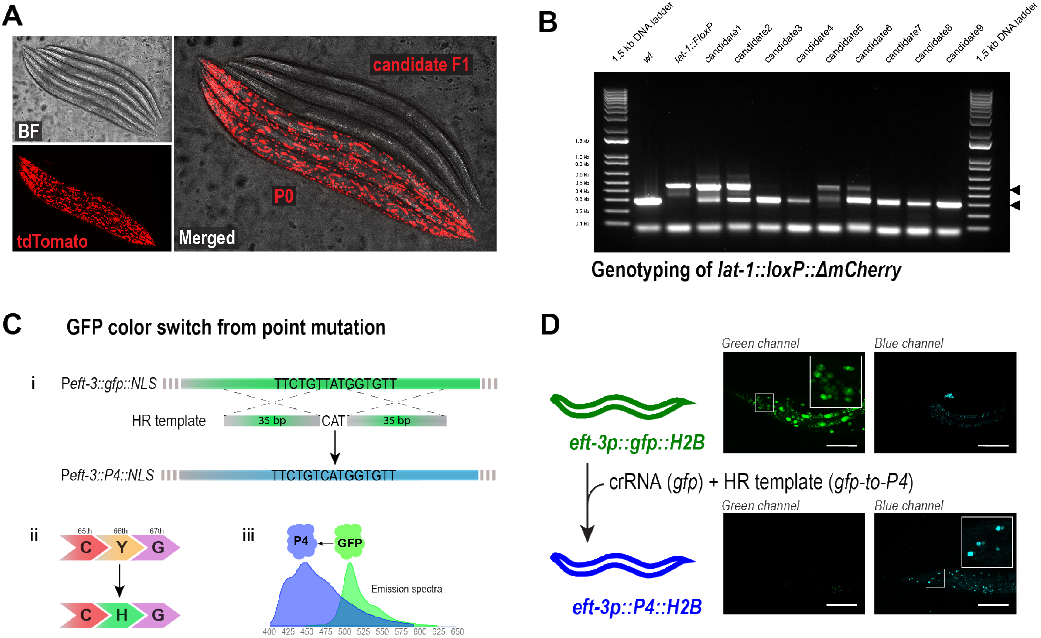
FLInt as co-CRISPR marker **A:** Representative images of a cohort of co-CRISPR’ed animals showing animals with the P0 phenotype and candidate F1. **B:** Screening PCR for *lat-1*::loxP. **C:** GFP to P4/BFP conversion as a homology-directed CRISPR marker. i) PAM sequence highlighted in bold, vertical line indicates Cas9 cutsite. ii) A single amino acid change in the GFP protein switches the absorption and emission wavelength to the blue. iii) Change in the emission spectrum from GFP to P4/BFP. **D:** Representative images of the co-converted GFP locus imaged for the GFP and BFP filtersets.

Together, our experiment revealed that the integration efficiency varies among the tdTomato inserted sites. The landing sites closer to the center appear to have higher efficiencies compared to the chromosome arms. Thus, in order to obtain the optimal integration efficiency, we suggest to use target loci closer to the center and in intergenic regions.

### Integrating existing extrachromosomal arrays into fluorescent safe habor loci

Lastly, we were interested in the targeted integration of existing extrachromosomal arrays into the tdTomato site without the use of mutagens such as UV or TMP that cause pleiotropic DNA defects and require subsequent outcrosses. To do so, we first crossed the target strain bearing the extrachromosomal array with the desired tdTomato marker strain, which we injected with the tdTomato CRISPR ingredients. We introduced, though, a slight variation following the observations in (Yoshina *et al*. 2016) in which they saw a correlation between integration frequency and fragmentation of the extrachromosomal array. This variation consisted of adding a crRNA that would target the Ampicillin resistance gene (which is present on the integrated vector plasmid), thus cutting the array in several pieces. Using the standard screening procedure (loss of NLS::tdTomato), we were able to recover 1 integrated line from 28 P0 (19 non-red F1) within 2-3 generations. The difference in the need of DNA cleavage between existing and *de novo* arrays probably lies in the fact that during the formation of the array, it already undergoes cleavage and assembly processes (Mello et al. 1991) that allow integration in one step. However, in a preexisting array there are no such events (Stinchcomb et al. 1985) and, thus, targeted editing facilitates NHEJ. With the present method, we demonstrated that a previously generated extrachromosomal array can be integrated into the tdTomato cleavage site without the drawbacks of random mutagenesis.

### Cas9-mediated disruption of tdTomato serves as a Co-CRISPR marker

A common bottleneck in the generation of CRISPR mutants is the efficient identification of successful gene edits. Without visible markers, PCR-based genotyping remains the ultimate option – a lengthy, tedious and potentially expensive process. Often, CRISPR-mediated genome editing in *C. elegans* is guided by a phenotypic conversion of an easily screenable co-CRISPR marker (Kim et al. 2014) that is eliminated after successful edits are isolated. In a successful edit, the mutated co-CRISPR locus results in an obvious phenotype which can be easily screened and distinguished from wildtype animals that were not edited. The marker phenotype thus, provides a visual representation of CRISPR efficiency and potentially reduces the number of progeny that eventually need to be sequenced to identify the desired edit. A large number of co-CRISPR marked progeny is indicative of putative edits at the gene of interest (GOI), always depending on the efficiency of the crRNA used for such locus. In *C. elegans* many co-CRISPR genes have been proposed. Among those, *pha-1, unc-22, sqt-1, unc-58, ben-1, zen-4* and *dpy-10* (Arribere et al. 2014; El Mouridi et al. 2017; Ward 2014; Dickinson et al. 2015; Kim et al. 2014) are popular, but may be problematic if its associated phenotype interferes with the GOI or is close to the target locus. Segregating alleles of genes that are in close proximity (e.g. *dpy-10* from other LGII genes) becomes problematic, since it depends on the genetic distance between the two genes. Likewise, co-CRISPR methods can result in subtle mutations at the co-CRISPR locus not phenotypically associated to it that can be confounded with the edit at the GOI (Rawsthorne-Manning *et al*. 2022).

We have already shown that the efficiency to induce Cas9-mediated double strand breaks of the crRNA for tdTomato is comparable, if not better, to the widely used *dpy-10* crRNA (Fig. 1C). Thus, tdTomato loci could pose an attractive alternative co-CRISPR locus, as its conversion does not result in any morphological or locomotion phenotype, and is ‘silent’. In addition, this could be beneficial when the co-CRISPR marker needs to be combined with a sublethal edit in essential genes that could lead to synthetic lethal phenotype (e.g. when combined with a *dpy-10* or *pha-1* co-CRISPR). Moreover, some phenotypic conversions (to a roller or a paralyzed animal), often preclude other phenotypic effects or can, in the worst cases, have a synthetic adverse effect with the desired gene modification. We specifically run into that problem when we designed a CRISPR edit for the GPCR *lat-1*, located physically close to *dpy-10*. We thus inserted a *loxP::*Δ*mCherry* site at the *lat-1* 3’ end and used the tdTomato in *oxTi390* IV as a coCRISPR marker. After injection into P0 animals, the jackpot plates contained non-red worms (Fig. 4A) as well as some dimmer/mosaic red F1 progenies, which we interpreted as excised in only one chromosome (see above). We only selected the non-red F1(s) for PCR screening of the *loxP::*Δ*mCherry* insertion at the *lat-1* locus (Fig. 4B) and successfully identified several candidates (edit efficiency = 28.38 *±* 13.25, n=7). Together, this result demonstrates that tdTomato can be used as a co-CRISPR locus that not only can be easily screened, but also does not interfere phenotypically with the target locus, what minimizes the need to unlink them.

Compared to *dpy-10(cn64)*, in which potentially successful edits can be identified as heterozygous repairs in *dpy-10* and segregate the GOI from the *dpy-10* locus, the excised tdTomato site is identified as homozygous. If elimination of the remains of tdTomato from the background is desired, the only possibility is outcrossing. However, the use of FLInt as co-CRISPR marker may involve more possibilities than for integration. Since only excision and not repair with the exogenous array is needed to successfully identify the canditates, the possible strains to be used increases with respect to the integration, in which we also need to account for higher probability of NHEJ repair with the array which. In addition, there is the possibility of using a tdTomato close to the GOI if the edit is difficult to screen in subsequent steps (e. g. a point mutation w/o visible phenotype in crosses). In those cases, PCR for the remaining tdTomato could be used in the screening processes. An important issue to consider is the choice of the tdTomato strain to use. Even though some of the 147 tdTomato target sites are mapped to genes, they do not result in visible phenotypes at 20C. However, whenever possible, intergenic safe habor sites should be used before starting an integration to avoid possible synthetic effects in downstream analyses.

### Single nucleotide conversion of GFP to BFP as a marker for HR-directed repair

Dominant co-CRISPR markers, as the widely employed *dpy-10(cn64)*, have the advantage that homology-directed repair can be visualized and distiguished from non homologous end joining repair directly in the F1 (Arribere *et al*. 2014). The use of tdTomato as a co-CRISPR marker, however, does not allow for such distinction in repair. To generate a co-CRISPR alternative for those cases, we took advantage of the possibility of changing the emission spectra of GFP from green to blue through a single nucleotide change (P4, Fig. 4C) (Heim et al. 1994). We designed a crRNA that cleaves the GFP sequence at the presumptive chromophore region together with an HR template that introduces the single point mutation to convert a tyrosine to an histidine at position 66. This genetic intervention switched the green emission spectrum of the GFP (508 nm) into the blue emission spectrum (448 nm). This simple modification can be made visible on a standard fluorescence microscope with an appropriate filter set (Fig. 4C (i, ii, iii)). After confirming that the crRNA efficiently cleaved the GFP sequence and led to a loss of GFP fluorescence in F1 animals, we added the HR template for the conversion and the corresponding mix for the GOI. We selected those F1 animals that showed a loss of GFP and emergence of blue fluorescence (Fig. 4D) which we used to screen for the edit at the GOI. However, because the P4/BFP fluorescence is rather weak as a heterozygous and might be difficult to see on standard epifluorescence stereoscope, similar strategies might provide larger contrast and easier screening. For example, the opposite conversion (from P4/BFP to GFP) yielded bright green fluorescence. Alternatively, a set of mutations centered around the tdTomato chromophore could be potentially mutated and the bright red turned into a bright green signal (Wiens et al. 2016). Together, these improvements might facilitate the use of GFP as a co-CRISPR marker, also when the GOI is linked closely to the traditional co-CRISPR locus and thus phenotypically interferes with the edits and/or cannot easily be unlinked through genetic breeding.

## Conclusion

In summary, we leveraged a library consisting of 147 marker strains that carry a single copy of a histone-tagged tdTomato and 142 strains with a nuclear localized GFP (Frøkjær-Jensen et al. 2014) as safe harbor landing sites for ectopic transgene integration. Importantly, all of these integrations reside at different locations on the six chromosomes in *C. elegans*, providing unprecedented flexibility in the genetic design and follow-up experiments. We demonstrated that these synthetic landing sites, encoding an ubiquitously expressed fluorescent protein, aided the identification of successful edits during transgene integration with several significant advantages: first, the integration site is known and precisely mapped, second, screening is facilitated through interference with the bright fluorescent signal indicating successful integration, third, a single crRNA can be used for all tdTomato landing sites, and, finally, because the intergenic landing site is known, the transgene integration does not cause any inadvertent phenotypes and defects. Together, these improvements in single shot transgenesis greatly reduce the time needed to screen for stable mutants, is flexible and cost effective, and has the potential to greatly accelerate research in *C. elegans*. In principle, this method can be extended to other invertebrate, vertebrate and mammalian model systems in which a single copy fluorescent gene is available as gene editing target sites.

## Data availability

All strains and plasmids generated during this study are available upon request to the corresponding author. Strains harboring the safe landing site are available through CGC and their information is accessible on wormbuilder.org.

## Acknowledgments

We would like to thank the NMSB lab for trouble shooting and beta-testing the procedure, Ravi Das for discussions and Julian Ceron for comments on the procedures and the manuscript, and the CGC (National Institutes of Health - Office of Research Infrastructure Programs (P40 OD010440)) for providing research reagents.

## Author contribution

NM and MK conceived the project, NM and MP performed experiments and analyses, MK and MP supervised the project and MK contracted funding. All authors wrote the first draft.

## Supplementary Materials

**Supplementary Figure S1**

**Supplementary Figure S2**

**Supplementary Figure S3**

**Supplementary Figure S4**

**Supplementary Figure S5**

## Supplementary Table

**Supplementary Table 1**

Detailed summary of the integration efficiency of all FLint loci tested with a standardized injection experiment in this study.

**Supplementary Table 2**

Summary and characteristics of strains generated using FLInt and other strains used in this study.

**Supplementary Table 3**

Plasmid(s) used and generated in this study.

**Supplementary Table 4**

DNA primer(s)

**Supplementary Table 5**

All crRNA(s) used in this study.

**Supplementary Table 6**

Homology directed repair templates.

**Supplementary Table 7**

tdTomato CRISPR mix preparation

**Supp. Fig.1.**
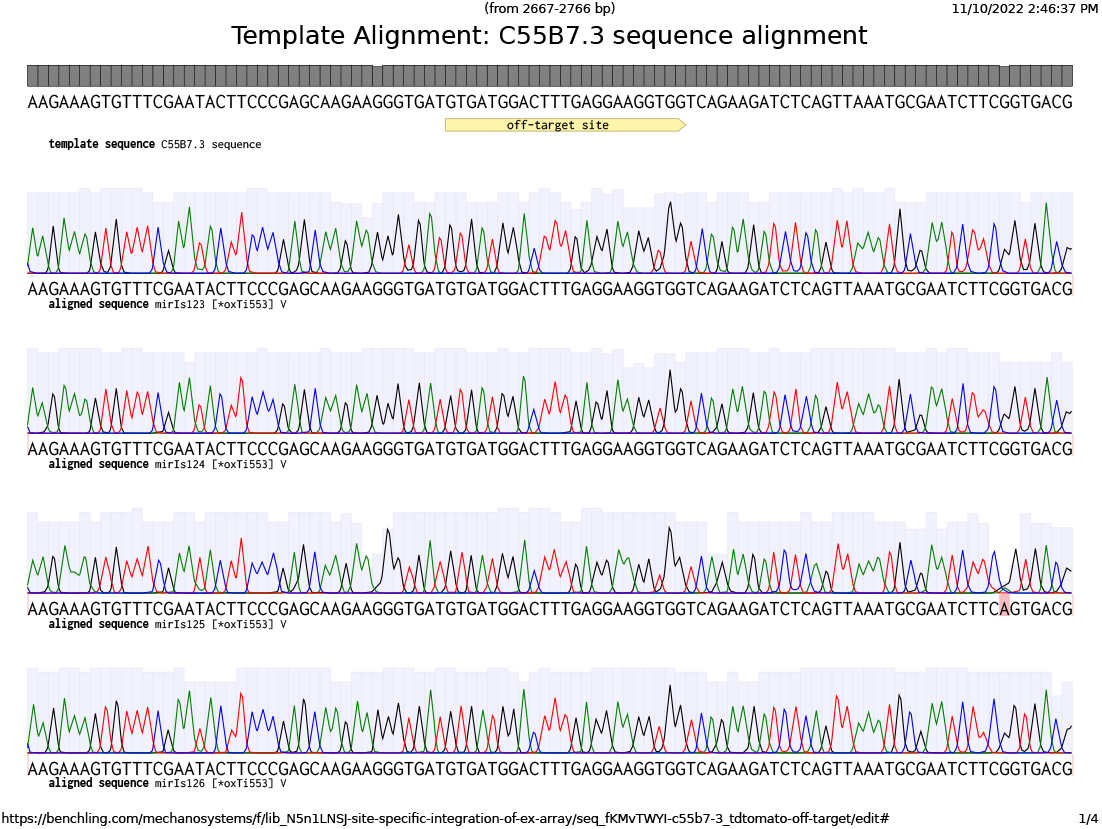
Sequencing results for predicted off-target site in C55B7.3 The employed tdTomato crRNAs are predicted to recognize the C55B7.3 sequence with four mismatched bases. However, no sequence defects have been detected in a total of 50 edited strains, 9 of which are shown here.

**Supp. Fig.2.**
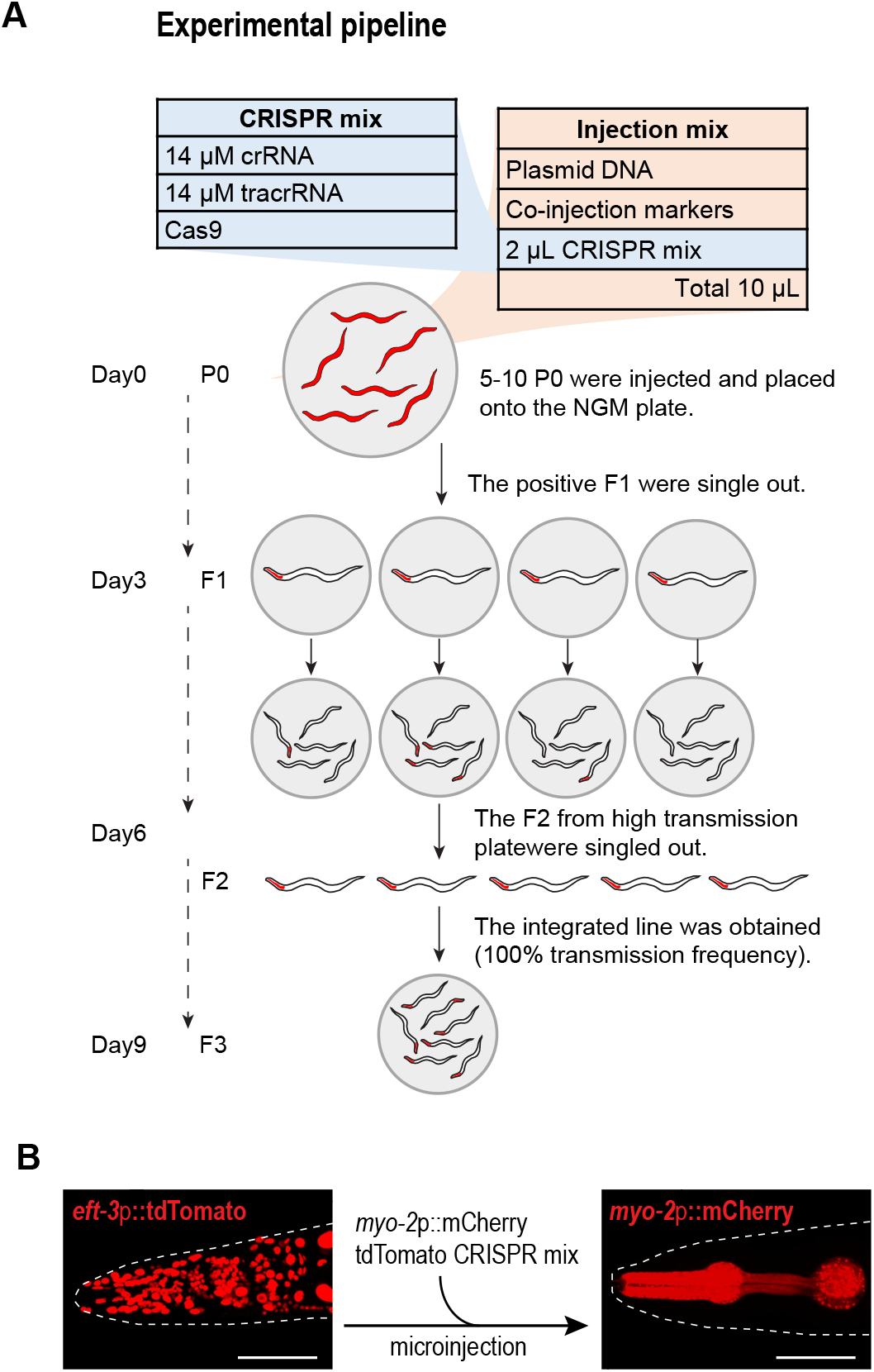
Experimental pipeline for FLInt integrations **A:** General procedure for transgene integration into a tdTomato locus. **B:** Representative photograph of an animal before and after successful integration. The correlation of a loss in red nuclear signal together with transgene fluorescence indicates successful integration.

**Supp. Fig.3.**
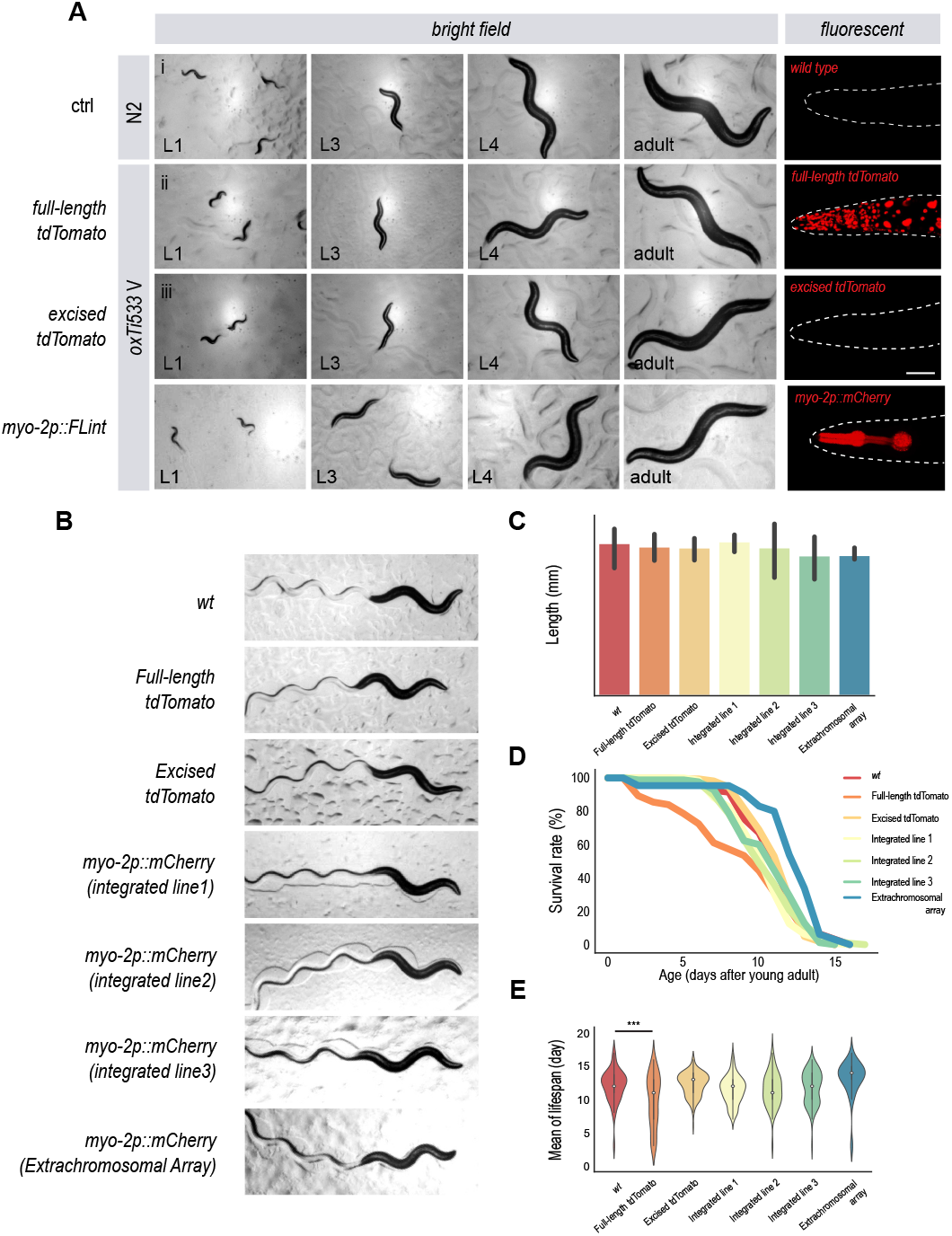
Health and lifespan of FLInt integrants **A:** Developmental time course of (i) wt animals, (ii) original strain with landing site (iii) recombined animals with no integration and (iv) FLInt animals from L1-adult day 1 and their fluorescent signal. **B:** Tracks of individual genotypes used to demonstrate the FLInt strategy. **C:** Body length of the employed genotypes. N=3 independent replicates. All conditions are p*>*0.05, as tested with Anova, Tukey-corrected for multiple comparisons. **D:** Lifespan curve of the FLInt animals. **E:** Lifespan distribution of the FLInt animals. p-values derived from a two sided Anova, Tukey-corrected test for multiple comparisons.

**Supp. Fig.4.**
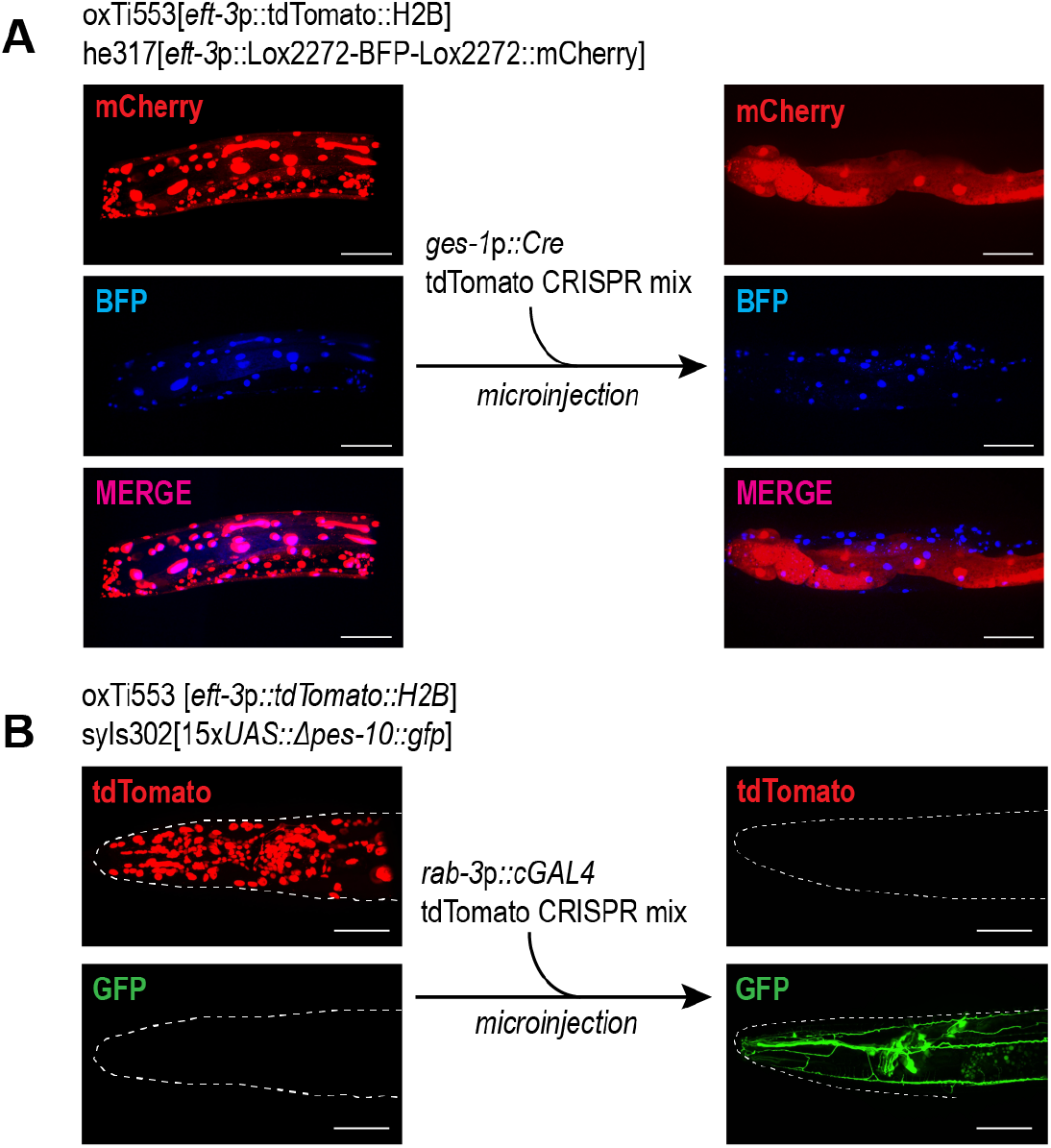
Using tdTomato FLInt as a co-transformation marker in drive/effector binary systems **A:** Integration of a recombinase enzyme (*ges-1*p::CRE) without need of a co-injection marker in *loxP* recombination marker background and screening by the BFP-to-mCherry color switch in CRE-expressing tissue (intestine). **B:** Integration of a transcription factor (*rab-3*p::cGAL4) in *UAS::gfp* background strain screening the GFP expression in *C. elegans* nervous system to isolate positive events.

**Supp. Fig.5.**
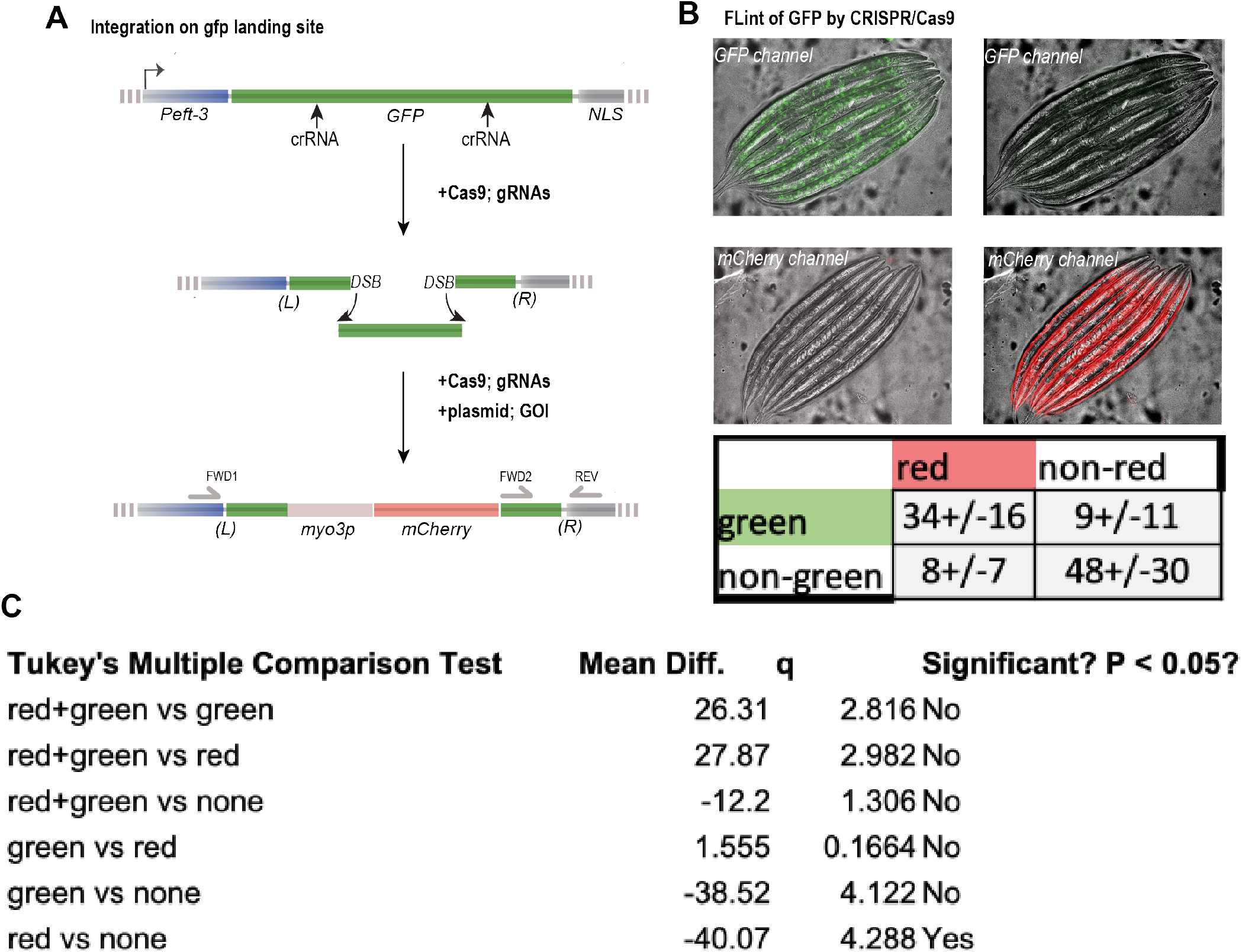
FLInt of GFP as a target site **A:** Schematic of the single copy loci and the gene replacement strategy. **B:** Representative outcome of the experiment is visualized by loss of GFP fluorescence and appearance of the transgene expression (BWM::mCherry). Below, table with number of observations (out of 100 animals; mean ± standard deviation) in tdTomato and GFP crRNA injected animals. **C:** Table with the Tukey-corrected ANOVA results for multiple comparisons of the outcome.

